# Blood-derived mitochondrial DNA copy number is associated with gene expression across multiple tissues and is predictive for incident neurodegenerative disease

**DOI:** 10.1101/2020.07.17.209023

**Authors:** Stephanie Y. Yang, Christina A. Castellani, Ryan J. Longchamps, Vamsee K. Pillalamarri, Brian O’Rourke, Eliseo Guallar, Dan E. Arking

## Abstract

**Background:** Mitochondrial DNA copy number (mtDNA-CN) can be used as a proxy for mitochondrial function and is associated with a number of aging-related diseases. However, it is unclear how mtDNA-CN measured in blood can reflect risk for diseases that primarily manifest in other tissues. Using the Genotype-Tissue Expression Project, we interrogated the relationships between mtDNA-CN measured in whole blood and gene expression from whole blood as well as 47 additional tissues.

**Results:** We evaluated associations between blood-derived mtDNA-CN and gene expression in whole blood for 418 individuals, correcting for known confounders and surrogate variables derived from RNA-sequencing. Using a permutation-derived cutoff (p<2.70e-6), mtDNA-CN was significantly associated with expression for 721 genes in whole blood, including nuclear genes that are required for mitochondrial DNA replication. Significantly enriched pathways included splicing (p=1.03e-8) and ubiquitin-mediated proteolysis (p=2.4e-10). Genes with target sequences for the mitochondrial transcription factor NRF1 were also enriched (p=1.76e-35).

In non-blood tissues, there were more significantly associated genes than expected in 30 out of 47 tested tissues, suggesting that global gene expression in those tissues is correlated with mtDNA-CN. Pathways that were associated in multiple tissues included RNA-binding, catalysis, and neurodegenerative disease. We evaluated the association between mtDNA-CN and incident neurodegenerative disease in an independent dataset, the UK Biobank, using a Cox proportional-hazards model. Higher mtDNA-CN was significantly associated with lower risk for incident neurodegenerative disease (HR=0.73, 95% CI= 0.66;0.90).

**Conclusions:** The observation that mtDNA-CN measured in whole blood is associated with gene expression in other tissues suggests that blood-derived mtDNA-CN can reflect metabolic health across multiple tissues. Key pathways in maintaining cellular homeostasis, including splicing, RNA binding, and catalytic genes were significantly associated with mtDNA-CN, reinforcing the importance of mitochondria in aging-related disease. As a specific example, genes involved in neurodegenerative disease were significantly enriched in multiple tissues. This finding, validated in a large independent cohort study showing an inverse association between mtDNA-CN and neurodegenerative disease, solidifies the link between blood-derived mtDNA-CN, altered gene expression in both blood and non-blood tissues, and aging-related disease.

## BACKGROUND

Mitochondria perform multiple essential metabolic functions including energy production, lipid metabolism, and signaling for apoptosis. Mitochondria possess circular genomes (mtDNA) that are distinct from the nuclear genome. While cells typically only possess two copies of the nuclear genome, they contain 100s to 1000s of mitochondria, and each individual mitochondrion can hold 2-10 copies of mtDNA resulting in wide variation in mtDNA copy number (mtDNA-CN) [1]. The amount of mtDNA-CN also varies widely across cell types, with higher energy demand cell types typically possessing higher levels of mtDNA-CN [1–3]. Due to the importance of mitochondria in metabolism and energy production, mitochondrial dysfunction plays a role in the etiology of many human diseases [4]. mtDNA-CN has been shown to be a proxy for mitochondrial function, and is consequently an attractive biomarker due to its ease of measurement [5,6]. Indeed, low levels of mtDNA-CN in peripheral blood have been associated with an increased risk for a number of chronic aging-related diseases including frailty, kidney disease, cardiovascular disease, heart failure, and overall mortality [7–10].

Crosstalk between the mitochondrial and nuclear genomes is essential for maintaining cellular homeostasis. Many essential mitochondrial proteins are encoded by the nuclear genome, and expression of these nuclear genes must be modified to match mitochondrial activity. Likewise, mitochondrial activity must respond to cellular energy demands. Polymorphisms in the nuclear genome have been associated with changes in mitochondrial gene expression, and mitochondrial genome variation has been associated with changes in nuclear gene expression, suggesting interplay between the two genomes [11,12].

In cancer cells, mtDNA-CN alters gene expression through modifying DNA methylation [13,14]. Recent work from our lab has shown that mtDNA-CN is also associated with nuclear DNA methylation in noncancer settings [15]. Given that DNA methylation can modify gene expression, the current study seeks to explore the potential association between blood-derived mtDNA-CN and gene expression. Past work has shown that mtDNA-CN is associated with gene expression of nuclear-encoded genes in lymphoblast cell lines, but this may not reflect biological processes occurring in other tissues, especially after an extended culturing period [16]. Therefore, we leveraged data from the Genotype-Tissue Expression Project (GTEx), a cross-sectional study with gene expression data from multiple non-diseased postmortem tissues, to examine associations between mtDNA-CN and expression of both nuclear and mitochondrially-encoded genes [17]. We found that blood-derived mtDNA-CN was globally associated with increased gene expression in whole blood. Additionally, blood-derived mtDNA-CN was associated with gene expression in other, non-blood tissues across the body. Specifically, genes annotated with neurodegenerative disease pathways were significantly enriched, leading to a follow-up analysis that uncovered a novel association between blood-derived mtDNA-CN and incident neurodegenerative disease.

## RESULTS

### Determination and validation of mtDNA-CN metric

mtDNA-CN estimates were generated from whole genome sequences performed on DNA derived from whole blood using the ratio of mitochondrial reads to total aligned reads. As mtDNA-CN is known to be affected by cell type composition, cell counts for samples with available RNA-sequencing data were deconvoluted using gene expression measured in whole blood [18,19]. We identified a batch effect that resulted in significantly altered mtDNA-CN for individuals sequenced prior to January 2013. Therefore, only individuals sequenced after January 2013 were retained for analysis (Supp. Fig. 1). After quality control, outlier filtering, and normalization of the RNA-sequencing data, 418 individuals remained for analyses (see **Methods**).

To validate mtDNA-CN measurements in the filtered GTEx data, we determined the association between mtDNA-CN and known correlated measures, including age, sex, and neutrophil count [18,20,21]. We observed a significant association with neutrophil count (p=5e-05), with higher neutrophil count associated with lower mtDNA-CN. While not statistically significant, effect size estimates between mtDNA-CN and age (p=0.19) and sex (p=0.17) were also in the expected direction. Effect sizes estimates for age and neutrophils were also consistent with prior literature [22] (Supp. Table 1). Based on variance explained from previous studies, the current study was only powered to detect a significant effect for neutrophil count. For all downstream analyses, mtDNA-CN was the standardized residual from a linear regression model adjusted for age, sex, cell counts, ischemic time, and cohort (see **Methods**).

### Association of mtDNA-CN derived from whole blood with gene expression in blood

A priori, we expect that mitochondrially encoded gene expression would be positively correlated with mtDNA-CN. Likewise, multiple nuclear encoded genes are involved in the regulation of mtDNA replication, and thus, expression levels of these genes are expected to be correlated with mtDNA-CN [23,24]. We therefore evaluated the associations between mtDNA-CN and expression of these two classes of genes, correcting for cohort, sample ischemic time, genotyping PCs, age, race, and surrogate variables derived from RNA-sequencing data to capture known and hidden confounders (Supp. Fig. 2) [25].

To minimize the potential impact of outliers, we performed an inverse normal transformation on both the mtDNA-CN metric and the gene expression values. To evaluate the association between mtDNA-CN and mitochondrial RNA (mtRNA) levels, we used the median gene expression value calculated from scaled expression values across 36 mtDNA-encoded genes that passed expression thresholds (see **Methods**).

We observed a highly significant association between mtDNA-CN and overall mtRNA expression (p=9.10e-9) (Table 1), with 33 out of 36 individual mtDNA-encoded genes nominally significant (p<0.05) (Supp. Fig. 3).

**Table 1.**
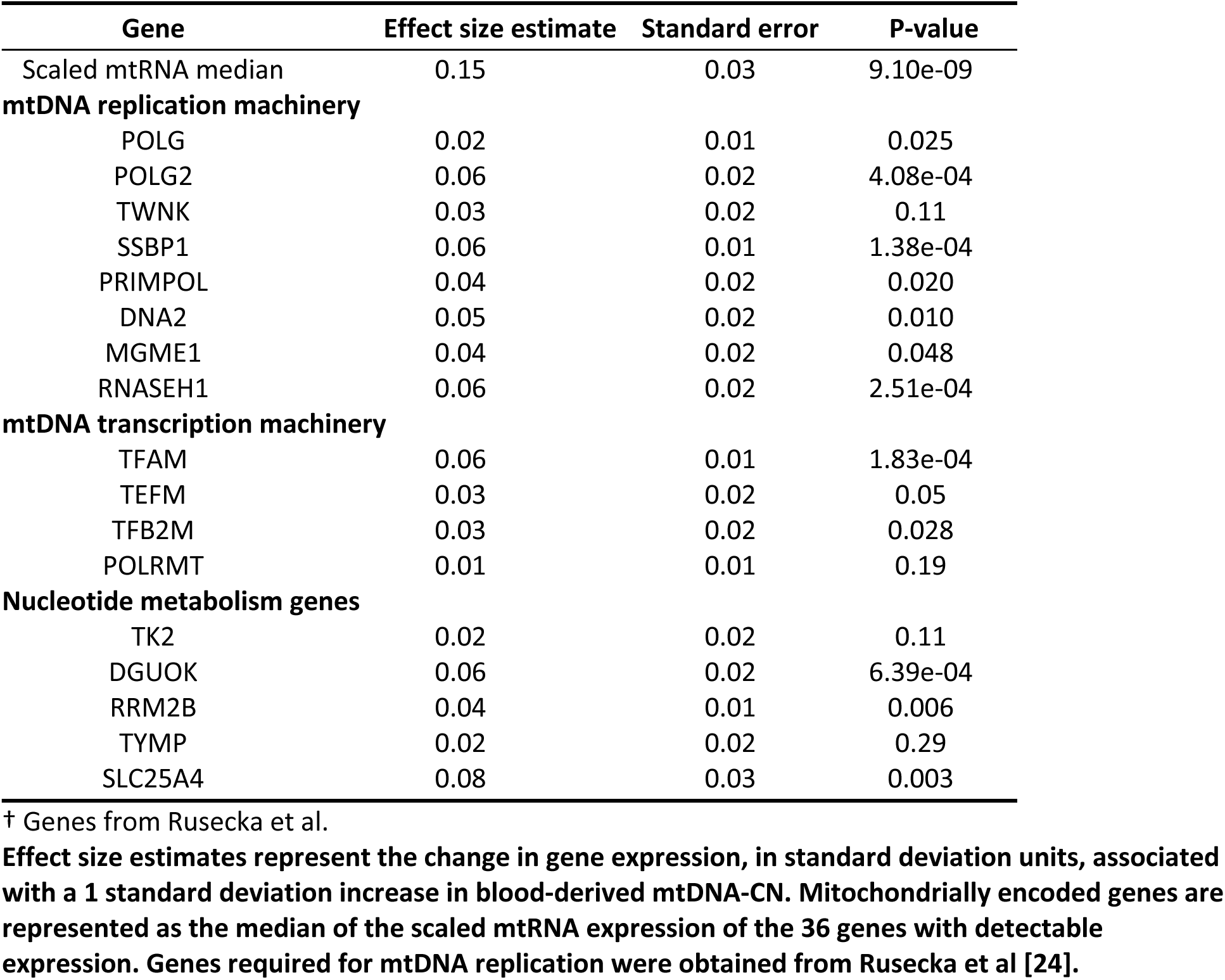
Blood-derived mtDNA-CN is positively associated with gene expression for mitochondrially encoded genes and nuclear encoded genes required for mtDNA replication.

| Gene | Effect size estimate | Standard error | P-value |
| --- | --- | --- | --- |
| Scaled mtRNA median | 0.15 | 0.03 | 9.10e-09 |
| **mtDNA replication machinery** |  |  |  |
| POLG | 0.02 | 0.01 | 0.025 |
| POLG2 | 0.06 | 0.02 | 4.08e-04 |
| TWINK | 0.03 | 0.02 | 0.11 |
| SSBP1 | 0.06 | 0.01 | 1.38e-04 |
| PRIMPOL | 0.04 | 0.02 | 0.020 |
| DNA2 | 0.05 | 0.02 | 0.010 |
| MGME1 | 0.04 | 0.02 | 0.048 |
| RNASEH1 | 0.06 | 0.02 | 2.51e-04 |
| **mtDNA transcription machinery** |  |  |  |
| TFAM | 0.06 | 0.01 | 1.83e-04 |
| TEFM | 0.03 | 0.02 | 0.05 |
| TFB2M | 0.03 | 0.02 | 0.028 |
| POLRMT | 0.01 | 0.01 | 0.19 |
| **Nucleotide metabolism genes** |  |  |  |
| TK2 | 0.02 | 0.02 | 0.11 |
| DGUOK | 0.06 | 0.02 | 6.39e-04 |
| RRM2B | 0.04 | 0.01 | 0.006 |
| TYMP | 0.02 | 0.02 | 0.29 |
| SLC25A4 | 0.08 | 0.03 | 0.003 |

† Genes from Rusecka et al.
**Effect size estimates represent the change in gene expression, in standard deviation units, associated with a 1 standard deviation increase in blood-derived mtDNA-CN. Mitochondrially encoded genes are represented as the median of the scaled mtRNA expression of the 36 genes with detectable expression. Genes required for mtDNA replication were obtained from Rusecka et al [24].**

In addition to genes coding directly for mtDNA replication machinery, genes involved in mtDNA transcription and nucleotide metabolism are also required for mtDNA replication. The mtDNA transcription machinery provides the RNA primers used in mtDNA replication and nucleotides are needed to synthesize new mtDNA molecules. Of the 17 mtDNA major replication genes tested [24], all were positively associated with mtDNA-CN, as would be expected based on gene function; 8 of them were nominally significant (p<0.05), and were 4 significant after Bonferroni correction (p<2.94e-3) for multiple testing (Table 1).

To identify additional genes and pathways associated with mtDNA-CN, we performed a transcriptome-wide analysis. There was an overall inflation of test statistics, which we quantified using the genomic inflation factor (lambda = 4.71) [26]. Two-stage permutation testing demonstrated no inflation in null datasets, suggesting that this inflation represents a true global association between blood-derived mtDNA-CN and gene expression (Supp. Fig. 4).

When stratified by gene functional categories [27], all categories showed elevated test statistics, but protein-coding genes were the most enriched (lambda=7.44) (Fig. 1). Gene expression levels of most of the nominally significant genes were positively correlated with mtDNA-CN (7769 genes with positive t-values vs. 285 genes with negative t-values). While much of this positive skewing is due to correlated gene expression, permuted datasets demonstrate that this positive shift is significant (P<0.001, Supp. Fig. 5), perhaps reflecting a more active transcriptional state associated with higher mtDNA-CN. Only two negatively associated genes passed permutation cutoff (p=2.7e-6), *CAMP* (p=1.58e-8) and *PGLYRP1* (p=1.78e-7), both of which are involved in innate immunity.

**Figure 1.**
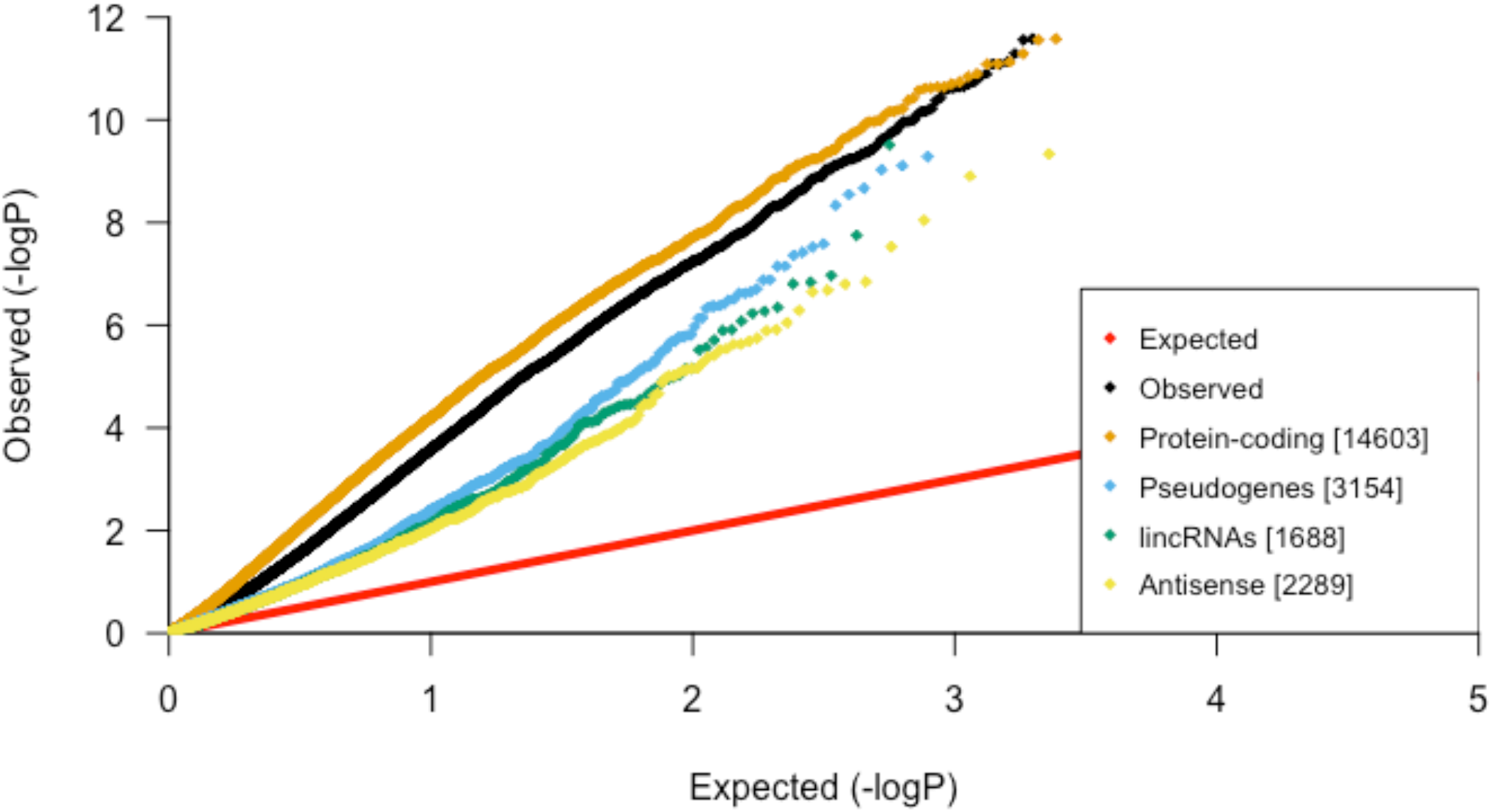
Global inflation of test statistics from linear regressions between blood-derived mtDNA-CN and gene expression in blood. After stratification by gene category, protein-coding genes have the most inflation, suggesting that mtDNA-CN is strongly associated with genes that code for proteins.

### Gene set enrichment analysis uncovers gene regulatory networks in whole blood

To identify specific molecular pathways, transcription factors, and gene ontologies associated with mtDNA-CN in whole blood, we performed gene set enrichment analyses [28] using gene sets obtained from the Molecular Signatures database (MSigDB) [29–33]. Previous studies have shown that cross-mappability can lead to false pseudogene positives in eQTL association studies [34]; we therefore excluded pseudogenes from subsequent analyses. Significantly associated KEGG pathways included “Spliceosome” (p=1.03e-8) and “Ubiquitin-mediated proteolysis” (p=2.4e-10) (Table 2).

**Table 2.**
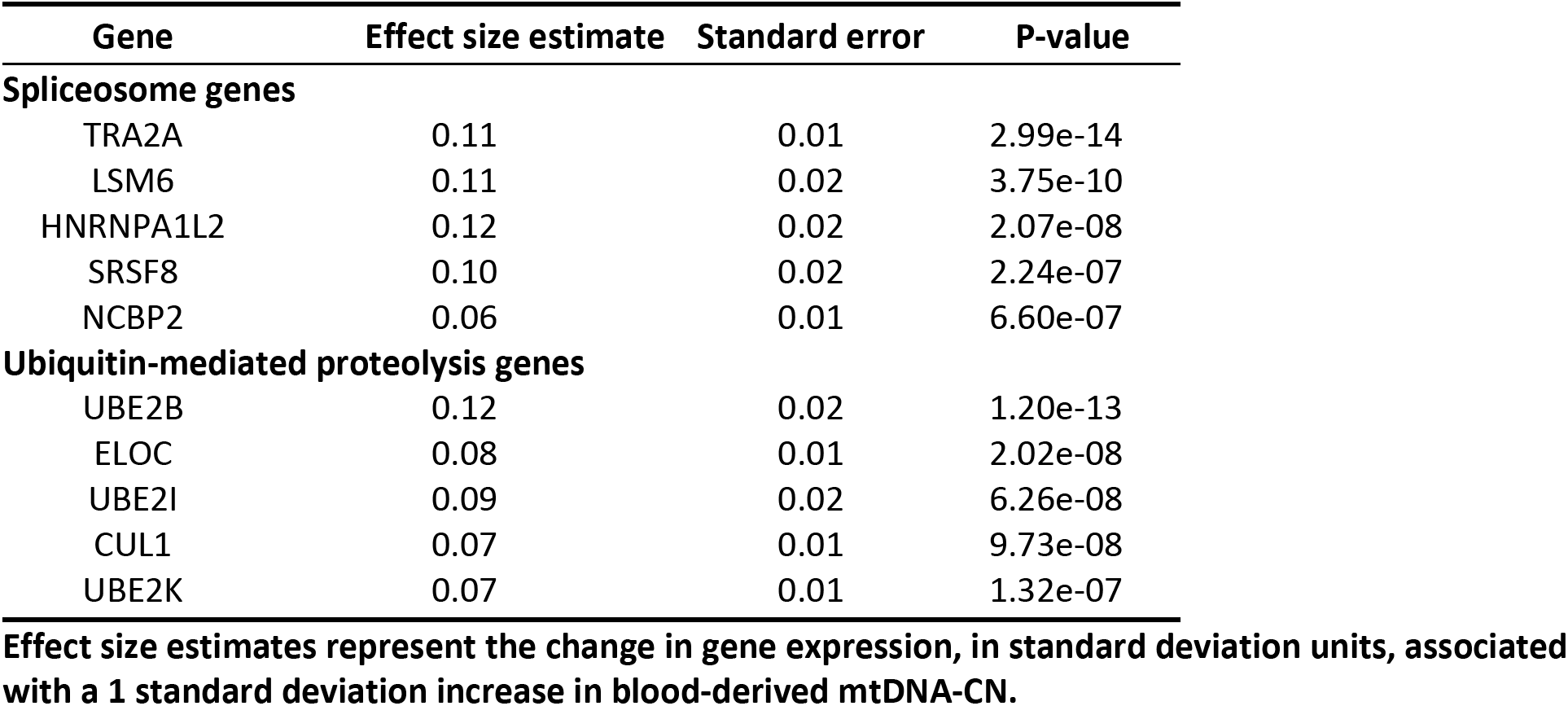
Top 5 genes that were most significantly associated with mtDNA-CN within the “Spliceosome” and “Ubiquitin-mediated proteolysis” KEGG pathways.

A number of transcription factor target sequences were also significantly enriched, including those for ELK1 (p=8.58e-66), NRF1 (p=1.76e-35), GABPB (p=3.54e-21), YY1 (p=3.14e-19), and E4F1 (p=3.98e-15). All of these transcription factors regulate genes that play a role in mitochondrial function [35–39]. Gene expression levels of these transcription factors were all positively correlated with mtDNA-CN, with 5 out of 6 nominally significant, and 3 remaining significant after Bonferroni correction (p<8.33e-3) (Table 3).

**Table 3.** Transcription factors whose targets are enriched for association with blood-derived mtDNA-CN are nearly all nominally significantly associated with blood-derived mtDNA-CN.

| Gene | Effect size estimate (gene expression) | Standard error (gene expression) | P-value (gene expression) | P-value (enriched target sequences) |
| --- | --- | --- | --- | --- |
| NRF1 | 0.03 | 0.02 | 0.07 | 1.76e-35 |
| YY1 | 0.07 | 0.01 | 1.78e-06 | 3.14e-19 |

| Gene | Effect size estimate<br>(gene expression) | Standard error<br>(gene expression) | P-value<br>(gene expression) | P-value<br>(enriched target sequences) |
| --- | --- | --- | --- | --- |
| GABPB2 | 0.09 | 0.02 | 1.51e-09 | 3.54e-21 |
| GABPB1 | 0.03 | 0.01 | 0.048 | 3.54e-21 |
| E4F1 | 0.05 | 0.01 | 2.01e-04 | 3.98e-15 |
| ELK1 | 0.04 | 0.02 | 0.021 | 8.58e-66 |

**Effect size estimates, standard errors, and p-values from a linear regression between transcription factor gene expression and blood-derived mtDNA-CN. Transcription factors shown are those whose targets were significantly enriched for association with blood-derived mtDNA-CN.**

Many mitochondrially-related cellular component gene ontology (GO) terms were significant, including “Mitochondrion” (p=7.77e-23), “Mitochondrial part” (p=2.79e-15), and “Mitochondrion organization” (p=2.87e-14) (Fig. 2) [40]. Additional significantly associated GO terms included “ubiquitin ligase complex” (p=6.6e-18) and “spliceosomal complex” (p=4.46e-14), supporting the KEGG pathway findings. Genes with substantial evidence of mitochondrial localization, determined through integration of several genome-scale datasets, were obtained from MitoCarta2.0 and demonstrated significant enrichment (p=8.22e-21) [41].

**Figure 2.**
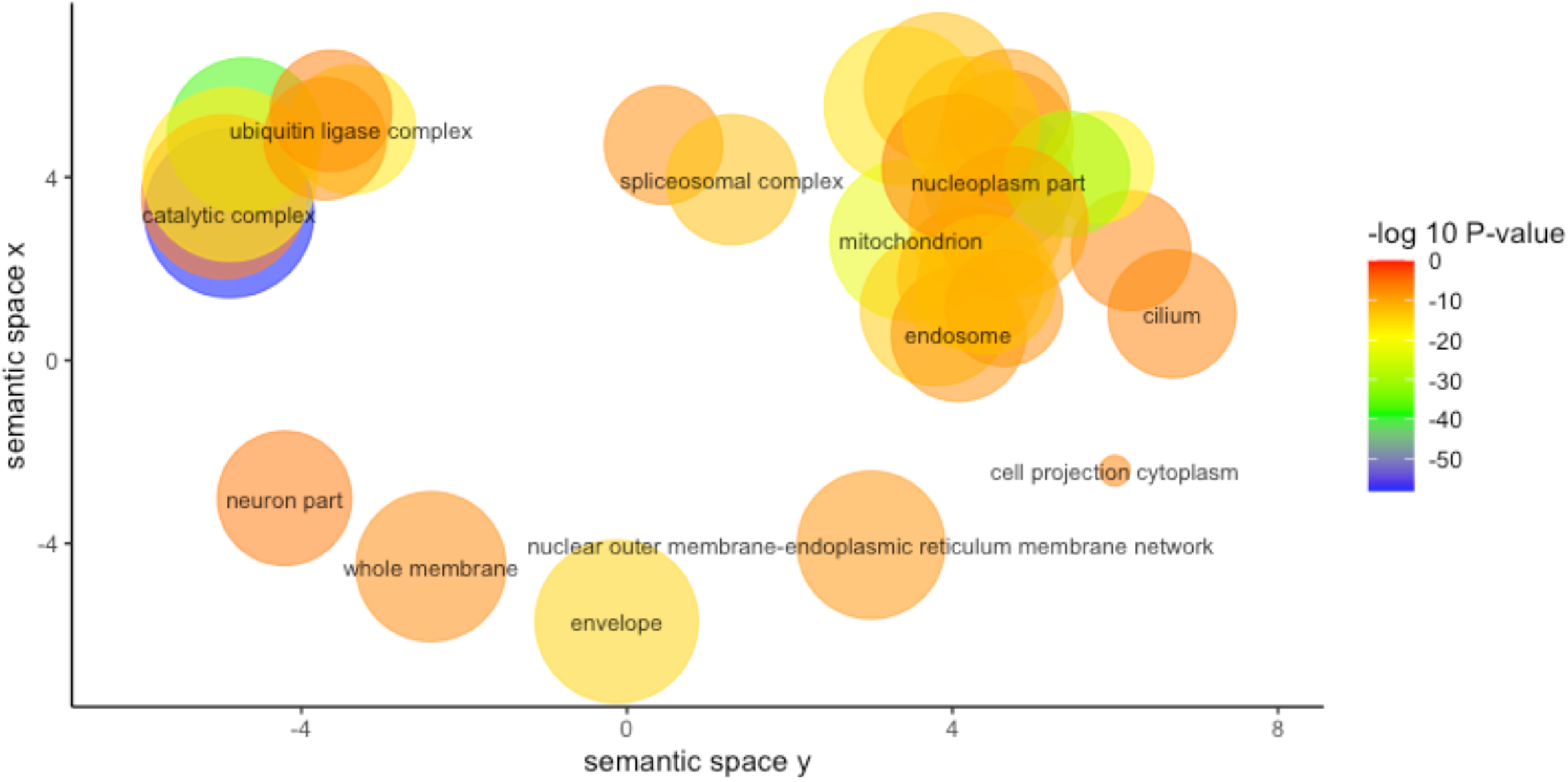
REVIGO visualization of GO Cellular Component terms significantly associated with mtDNA-CN after removal of redundant GO terms. Size of the circle represents the relative number of genes in each gene set, color represents significance. Axis represent semantic similarities between GO terms; GO terms that are more similar will cluster with one another.

### Cross-tissue analysis reveals associations between gene expression in multiple tissues and blood-derived mtDNA-CN

mtDNA-CN measured in blood has been associated with a number of aging-related diseases including chronic kidney disease, heart failure, and diabetes [10,42,43]. Given that these diseases primarily manifest in non-blood tissues, we evaluated associations between blood-derived mtDNA-CN and gene expression measured from 47 additional tissues that had greater than 50 samples after filtering. Though blood-derived mtDNA-CN appears to be associated with gene expression in other tissues, we did not observe a significant association between blood-derived mtDNA-CN and scaled mtRNA gene expression in any tissue other than blood, and only 2 out of 47 tested tissues had nominally significant associations between tissue-specific scaled mtRNA expression and blood-derived mtDNA-CN (Uterus [p = 0.004], Heart – Left Ventricle [p = 0.017]). (Supp. Table 2). However, mtRNA expression for 35/47 non-blood tissues was positively associated with blood mtDNA-CN, which is more than what would be expected by chance (p<0.001). This suggests that while our study may be underpowered to detect a significant association in individual tissues due to small sample sizes, mtDNA-CN measured in blood is broadly correlated with mtDNA-CN in other tissues.

We calculated genomic inflation factors for each tissue to quantify test statistic inflation. Genomic inflation factors were elevated across multiple non-blood tissues, suggesting that blood-derived mtDNA-CN was broadly associated with gene expression in other tissues (Fig. 3). To determine true signal from noise, we performed 100 two-stage permutations for each tissue and obtained a genomic inflation factor lambda cutoff of >1.20 representing a significant elevation of lambda (study-wide p<0.05). Using this cutoff, we identified 30 non-blood tissues with a global inflation of test statistics [Supp. Table 3].

**Figure 3.**
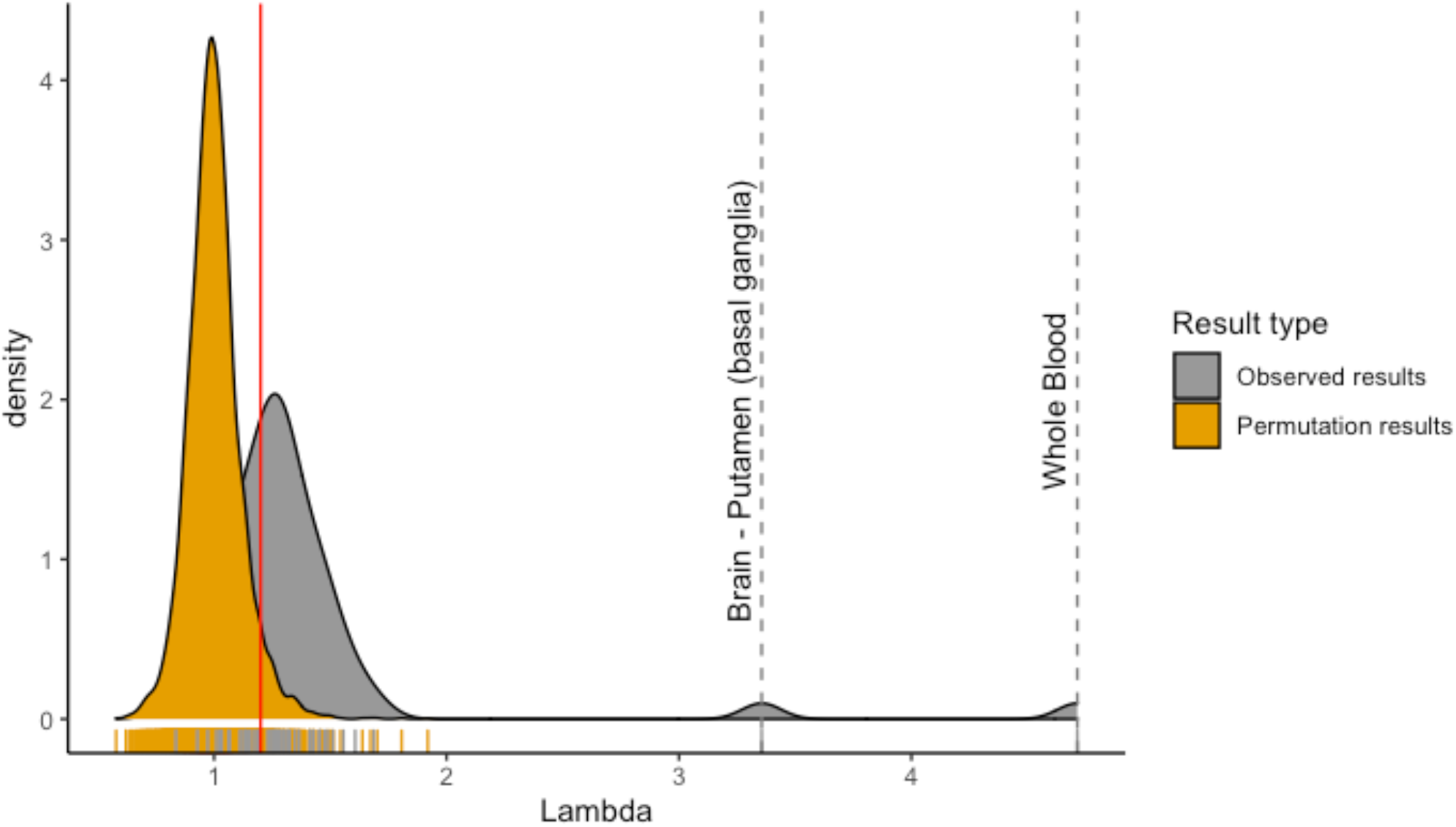
Observed genomic inflation factors are significantly different from permuted genomic inflation factors for certain tissues. Higher genomic inflation factor represents increased global associations between blood-derived mtDNA-CN and gene expression in a specific tissue. Permuted genomic inflation factors were obtained using two-stage permutation testing.

Other than blood, the most strongly enriched tissue was the putamen region of the brain, with a lambda of 3.27. We note that the two cell lines, EBV transformed lymphocytes (lambda=0.84) and cultured fibroblasts (lambda=0.84), showed no global inflation of test statistics, suggesting that blood-derived mtDNA-CN loses its association with gene expression after the cell-culturing process.

To examine the similarity of associations of mtDNA-CN observed in blood with other tissues, we calculated Spearman’s rank correlation coefficients between effect estimates for blood-derived mtDNA-CN on blood gene expression (β_blood_) and effect estimates for blood-derived mtDNA-CN on gene expression in other tissues (β_tissue_). All genes that passed a permutation cutoff for significance in blood (p=2.7e-6, 721 genes total) were included. To distinguish tissues with correlations more extreme than baseline, we calculated reference correlations between blood and other tissues for randomly selected sets of genes. 26 tissues had observed values that were more extreme than the random gene sets (Supp. Table 4). Of these 26 tissues, 20 were among the 30 tissues with significantly inflated lambdas.

To identify pathways associated with mtDNA-CN across multiple tissues, we performed gene set enrichment analysis in each of the 30 tissues with a significant genomic inflation factor. Multiple terms were significant in greater than one tissue (Table 4), including terms related to oxidative phosphorylation and mitochondria, suggesting that mtDNA-CN derived from blood can reflect mitochondrial function occurring in other tissues.

**Table 4.**
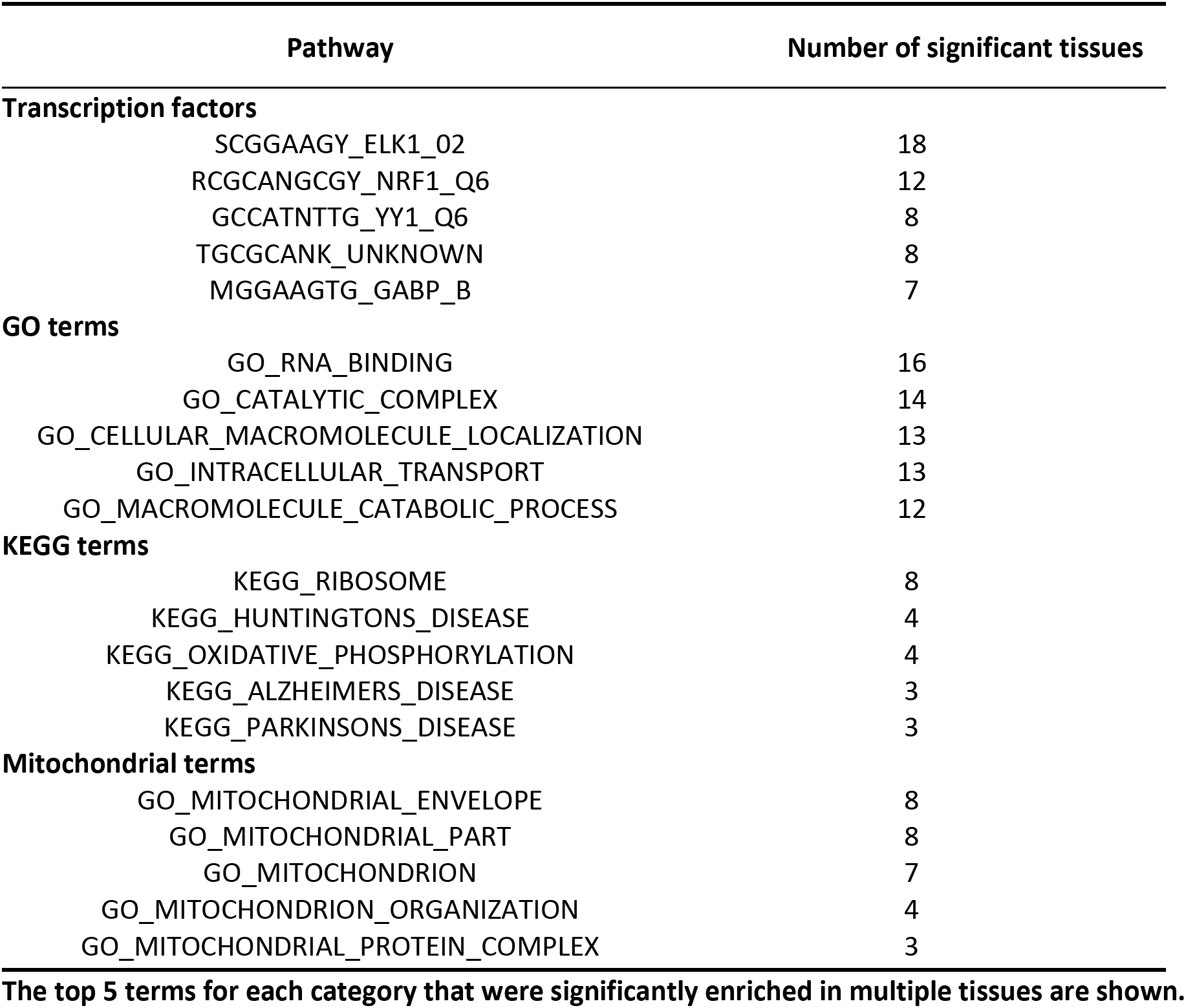
Pathways, transcription factor targets, and GO terms significantly enriched in multiple tissues.

### The top 5 terms for each category that were significantly enriched in multiple tissues are shown

ELK1 transcription factor binding sites were significantly enriched in 17 of the 30 significant tissues, and were also significant in whole blood, suggesting that mtDNA-CN may regulate ELK1 or vice versa. We note that gene expression for ELK1 was nominally significantly associated (p<0.05) with blood-derived mtDNA-CN in 12 of the 18 tissues for which ELK1 targets were significantly enriched (Supp. Fig. 6). Effect estimates for ELK1 targets were generally consistent with the directionality of ELK1 effect estimates. For example, in blood, where ELK1 expression is positively associated with mtDNA-CN, 747/750 (99.6%) nominally significant ELK1 target genes were positively associated. On the other hand, mtDNA-CN was negatively associated with nerve ELK1 gene expression, and 204/306 (66.67%) nominally significant ELK1 target genes were also negatively associated. Of note, nearly all the noted transcription factors were ubiquitously expressed throughout the body, except for ELK1, which is not expressed in Brain Putamen or Spinal Cord (Supp. Fig. 7).

To identify genes driving enrichment of significant pathways in multiple tissues, we performed a random effects meta-analysis for all expressed genes using effect size estimates from all 47 non-blood tissues. Strikingly, genes encoding both the large and small ribosomal subunits were negatively associated with blood-derived mtDNA-CN across all tested tissues, implying an inverse relationship between ribosomal abundance and mitochondrial DNA quantity (Table 5).

**Table 5.** Random-effects meta-analysis for genes driving the enrichment of pathways in multiple tissues.

| Gene | Meta effect size estimate | Meta standard error | Meta p-value |
| --- | --- | --- | --- |
| **ELK1 targets** |  |  |  |
| STARD3 | 0.05 | 0.00 | 1.49e-17 |
| EIF5A | 0.08 | 0.01 | 4.97e-17 |
| ERH | 0.07 | 0.01 | 8.82e-17 |
| **RNA-binding genes** |  |  |  |
| SUZ12 | 0.06 | 0.00 | 4.49e-18 |
| C1D | 0.11 | 0.01 | 8.03e-18 |
| MRPL23 | -0.09 | 0.01 | 8.37e-18 |
| **Ribosome genes** |  |  |  |
| RPL34 | -0.07 | 0.01 | 8.57e-16 |
| RPS27 | -0.08 | 0.01 | 1.35e-15 |
| RPL39 | -0.08 | 0.01 | 1.16e-14 |
| **Mitochondrial part genes** |  |  |  |
| MRPL23 | -0.09 | 0.01 | 8.37e-18 |
| MTERF3 | -0.06 | 0.00 | 3.12e-17 |
| MICU3 | 0.09 | 0.01 | 3.34e-17 |

### Meta-analysis results are from all 47 tested tissues, excluding effects from whole blood. Top 3 most significant genes for each pathway are shown

Huntington’s disease (HD), Parkinson’s disease (PD), and Alzheimer’s disease (AD) were among the most significantly associated KEGG pathways that appear in multiple tissues (Table 5). This is an intriguing finding, given the known role of mitochondria in neurodegenerative disease [44].

While neurodegenerative disorders primarily manifest in nervous tissues [45], we observed significant enrichment of disease pathways in colon, pancreas, and testis tissues. When limiting our query to brain tissues, HD and PD were nominally significantly enriched in cerebellum, caudate (basal ganglia), and cortex, while AD was nominally significantly enriched in cerebellum and spinal cord (Supp. Fig 8).

### mtDNA-CN is associated with incident neurodegenerative disease in the UKBiobank

To examine the association between mtDNA-CN and neurodegenerative disease risk, we used the UK Biobank (UKB) [46], a prospective cohort study with whole exome sequencing for ~50,000 individuals. mtDNA-CN was derived from whole exome sequencing and adjusted for sequencing artifacts, age, and sex. Analysis was restricted to individuals of European descent, and individuals with blood cell type count outliers were excluded. Using a Cox proportional-hazards model and adjusting for age and sex, we evaluated the relative risk of neurodegenerative disease associated with mtDNA-CN. Median follow-up time was approximately 10 years. Although the number of incident events was small, mtDNA-CN was significantly associated with Parkinson’s disease (HR=0.75, CI=0.60;0.99) and Alzheimer’s disease (HR=0.59, CI=0.44;0.81), and approached significance for non-Alzheimer’s dementia (HR=0.81, CI=0.65;1.02). Consistent with other aging-related diseases [7,9], higher mtDNA-CN was associated with lower risk for developing incident neurodegenerative disease (Table 6). A combined analysis for all individuals with incident neurodegenerative disease revealed a consistent strongly significant association (HR=0.73, CI=0.66;0.90).

**Table 6.** mtDNA-CN is associated with incident neurodegenerative disease.

| Disease | Hazard ratio | Confidence interval | Number of cases/controls | P-value |
| --- | --- | --- | --- | --- |
| Parkinson's disease | 0.75 | 0.60;0.99 | 63/39,044 | 0.030 |
| Alzheimer's disease | 0.59 | 0.44;0.81 | 41/39,100 | 0.001 |
| Dementia (excluding AD) | 0.81 | 0.65,1.02 | 74/39,048 | 0.074 |
| Combined neurodegenerative disease | 0.73 | 0.66;0.90 | 161/39,030 | 0.001 |

**Hazard ratios for incident neurodegenerative disease associate with a 1 standard deviation increase in whole blood mtDNA-CN estimated from Cox proportional-hazards models in the UKBiobank. Analysis was restricted to individuals of European descent, and individuals who were outliers for cell counts were excluded from analysis.**

## DISCUSSION

In this study, blood-derived mtDNA-CN was significantly associated with a host of blood-expressed genes. As expected, nearly all genes involved in mtDNA replication were significantly associated with mtDNA-CN in a positive direction. There was also a clear overall shift towards significant positive estimates, possibly indicating that increased mtDNA-CN is reflective of a more active transcriptional state. This finding is consistent with previous literature demonstrating that higher mitochondrial content is correlated with increased transcriptional activity [47,48]. Strikingly, the two negatively associated genes both play roles in innate immune function [41,42], suggesting that higher mtDNA-CN levels are correlated with decreased immune response. Mitochondria play a role in immune responses to pathogens in several ways; for example, mitochondrial DNA release from compromised mitochondria can trigger an intracellular antiviral response through the cGAS/STING pathway [51], binding of viral dsRNA to the mitochondrial antiviral signaling complex (MAVS) can trigger an interferon responses through STAT6 activation [52], and release of mitochondrial components from cells can bind to damage-associated molecular pattern (DAMP) receptors to trigger innate immune responses [53]. These novel findings correlating expression of mtDNA-CN with specific immune response genes in tissues represent an area for further investigation.

Gene set enrichment analyses revealed pathways potentially involved in mitochondrial DNA control, including ubiquitin-mediated proteolysis and splicing. Supporting this finding, Guantes et. al demonstrated that mitochondrial content modulates alternative splicing [47]. Additionally, we found that genes expressed in whole blood that were associated with blood-derived mtDNA-CN were enriched for target sequences for the ELK1, NRF1, YY1, GABPB, and E4F1 transcription factors. All of these transcription factors have been implicated in mitochondrial pathways, as ELK1 is associated with the mitochondrial permeability transition pore complex in neurons, NRF1 regulates expression of the mitochondrial translocase TOMM34, YY1 binds to and represses mitochondrial gene expression in skeletal muscle, GABPB is required for mitochondrial biogenesis, and E4F1 controls mitochondrial homeostasis [35–39]. Additionally, we found significant enrichment of signal for genes implicated in ubiquitin-mediated proteolysis and splicing. Given that mitochondrial quality control is regulated through ubiquitination, and that nuclear-encoded spliceosomes are involved in mtRNA splicing, our results likely implicate processes involved in mitochondrial DNA regulatory networks [54,55].

mtDNA-CN measured in one tissue has previously been found to be uncorrelated with mtDNA-CN in another tissue from the same individual [56]. We found that while mtRNA transcription in individual tissues was not significantly correlated with blood-derived mtDNA-CN, across all tissues, there was a significant enrichment for positive associations, suggesting a weak positive correlation between blood-derived mtDNA-CN and mtDNA-CN in other tissues. Moreover, we found that blood-derived mtDNA-CN was associated with various biological pathways in non-blood tissues (including mitochondrial function), providing a possible explanation as to why blood-derived mtDNA-CN is associated with aging-related diseases that primarily manifest in non-blood tissues. Further examination of pathways significant in multiple tissues revealed that ribosomal subunit genes were significantly negatively associated with mtDNA-CN. While there has been conflicting evidence on the relationship between mtDNA-CN and ribosomal content, our study revealed a strong inverse relationship between ribosomal DNA dosage and mtDNA-CN [16,47]. Importantly, since these are statistical associations, causal directionality cannot be determined between gene expression and blood-derived mtDNA-CN. Future follow-up studies are needed to determine functional causality for mtDNA-CN and gene expression.

Strikingly, KEGG pathways that were significantly enriched in multiple tissues included Huntington’s disease, Alzheimer’s disease, and Parkinson’s disease. These aging-related neurodegenerative diseases have all underlying mitochondrial pathologies [57–60] and dysregulated ubiquitination pathways [61]. In particular, mtDNA-CN has been implicated in Alzheimer’s disease [62–64] and cognitive function [65,66]. Further, the ELK1 transcription factor, whose target sequences were significantly enriched in 18 tissues, plays a role in multiple neurodegenerative diseases [67]. Finally, after finding that blood-derived mtDNA-CN was associated with expression of neurodegenerative disease genes, we used an independent dataset, the UK Biobank, and found that mtDNA-CN was significantly associated with incident neurodegenerative disease risk. Notably, the population of individuals used for the UK Biobank analyses is biased towards fewer smokers and fewer prevalent Parkinson’s events [68], which may impact the generalizability of our results. Despite this caveat, we show that blood-derived mtDNA-CN is significantly associated with gene expression from tissues across the body, and that higher mtDNA-CN is associated with decreased incident neurodegenerative disease risk.

## METHODS

### GTEx Sample acquisition

Whole genome sequences were downloaded from the GTEx version 8 cloud repository on 11/18/2020. RNA-sequencing data used for analyses was downloaded from the GTEx portal (http://gtexportal.org/home/datasets) on 06/18/2019 and phenotypes were obtained from dbGaP (phs000424.**v8**.p2).

### Estimation of mtDNA-CN

Samtools version 1.9 [69] was used to count the number of mitochondrial, unaligned, and total reads for each whole genome sequence. mtDNA-CN was estimated as the number of mitochondrial reads divided by the difference between the number of total reads and the number of unaligned reads to obtain a ratio of mtDNA to nuclear DNA. Whole genome is a highly accurate method for estimation of mtDNA-CN [22,56].

### Correcting mtDNA-CN for covariates

All statistical analyses were performed with R version 3.6.1. Cell type composition for whole blood samples was determined from RNA-sequencing using xCell [19], only allowing for deconvolution of cell types found in blood. A stepwise regression in both directions was used to select appropriate cell types to correct mtDNA-CN. To avoid model overfitting, correlated cell types (R>0.8) were removed. The final model used to adjust mtDNA-CN included neutrophils, hematopoietic stem cells, megakaryocytes, subject cohort, ischemic time, age, and sex. Power calculations were performed using R^2^ values from previous studies using the pwr package [22].

### Filtering pipeline

A batch effect due to sample collection and/or sequencing methods resulted in significantly altered mtDNA-CN for individuals who were sequenced prior to January 2013. To keep this from confounding the analysis, we excluded subjects with whole genome sequencing prior to January 2013 (Supp. Fig. 1). Individuals who had greater than 5×10^7^ unaligned whole genome sequence reads were also omitted from the analysis. Cell type outliers who were greater than 3 standard deviations (SDs) from the mean were excluded as well. Only one individual remained from the surgical cohort after filtering and therefore was also removed (Supp. Fig. 9).

### RNA-sequencing pipeline

GTEx version 8 RNA-sequencing data was downloaded from the GTEx website in transcripts per million (TPMs) and normalized using the trimmed mean of M-values method prior to analyses [70,71]. Genes with expression greater than 0.1 TPMs for at least 20% of samples were retained for analysis. To identify potential hidden confounders, we used surrogate variable analysis (SVA), protecting mtDNA-CN from SV generation [25]. SVs were associated with known covariates in the data, such as whether individuals were in the postmortem or the organ donor cohorts (Supp. Fig. 2). Individuals who were greater than 3 standard deviations from the mean for the first ten SVs were omitted from analysis. SV generation was performed iteratively 3 times.

### Linear model for evaluating associations

To reduce the influence of outliers, both the gene expression metric and the mtDNA-CN metric were inverse normal transformed prior to linear regression. We then tested for association using multiple linear regression, with mtDNA-CN as the predictor and gene expression as the outcome, correcting for SVs, sex, cohort, race, ischemic time, and the first three genotyping principal components.

### Genomic inflation factor calculation

Genomic inflation factors were calculated by squaring z-scores to obtain chi-squared values. The median observed chi-squared value was divided by the expected median to obtain lambda [26].

### Two-stage permutations

To determine an appropriate p-value cutoff, we created null datasets for permutation testing. First, a multiple linear regression model for the alternate hypothesis was used to obtain gene expression residuals. Second, a multiple linear regression model for the null hypothesis was used to obtain estimates for each gene. Residuals from the alternate model were than permuted and added to effect estimates from the null model to create null datasets. Permuted gene expression data was then tested for association with mtDNA-CN. Unless otherwise stated, permutations were performed 100 times. Minimum p-values from each permuted dataset were obtained, and the 5^th^ lowest p-value was utilized as a permutation cutoff.

### Annotation of gene categories

Gene annotations were downloaded from Gencode [27]. Test statistics were then stratified by gene type and observed and expected distributions were generated for each category.

### Overrepresentation of positive beta estimates

Percentage of positive effect estimates was calculated using all nominally significant genes in blood, dividing the number of nominally significant genes with positive effect estimates by the total number of nominally significant genes. Percentages for null distributions were calculated using 1000 permutations generated using the two-stage permutation method described above.

### Gene set enrichment analysis

To examine enrichment for genes in specific pathways, gene sets for KEGG pathways, transcription factor target sequences, and gene ontologies were downloaded from the Molecular Signatures database [25,27,49]. Then, using the absolute value of the t-scores from the regression model with mtDNA-CN, we performed a t-test of t-scores for genes in a specific pathway versus genes that were not contained in the pathway. Permutations using randomized t-scores were used to determine appropriate cutoffs for significance. To confirm that results were not driven by individual genes in a pathway with very large t-scores, we also performed t-tests using ranked t-scores as opposed to absolute value t-scores.

### REVIGO trimming and visualization of GO terms

For visualization of significantly enriched GO terms and elimination of redundant GO terms, REVIGO (http://revigo.irb.hr/) was used with the default settings except for the allowed similarity, which was set to medium (0.7) [40].

### Testing for associations between blood-derived mtDNA-CN and gene expression in other tissues

Filtering parameters and models for testing the association of blood-derived mtDNA-CN with gene expression in other tissues were identical to the pipeline used in whole blood. Only tissues with greater than 50 observations after filtering were tested. For tissues that had no variation in covariates, covariates were dropped from the linear model (i.e. sex was not used in the model for testing gene expression in reproductive organs and cohort was not used in the model for brain tissues).

### Spearman correlations for effect estimates with whole blood

All significant genes in whole blood that passed the permutation cutoff (p=2.7×10^−6^) were used for testing. Spearman correlations between effect estimates in blood and effect estimates in other tissues were calculated. To compare correlations for genes significant in blood with baseline correlation, we randomly selected 100 random genes and calculated correlations between blood estimates and specific tissue estimates for those genes. We repeated this random selection 100 times to generate multiple baseline correlation measures.

### Meta-analysis of genes driving specific ontologies

To calculate meta-analysis effect estimates and p-values, the R ‘*meta’* package [73] was used to perform a random-effects meta-analysis using all effect estimates and p-values for all tissues, excluding results from whole blood.

### Association of mtDNA-CN with neurodegenerative disease in UKB

Samtools version 1.9 was used to extract read summary statistics from 49,997 UK biobank whole exome sequences. An in-house perl script was then used to aggregate summary statistics into the number of total, mapped, unmapped, autosomal, chromosome X, chromosome Y, mitochondrial, random, unknown, decoy1, and decoy2 reads. 10-fold cross validation was used to select linear regression covariates to adjust the number of mitochondrial reads for. After correcting for these potential technical artifacts, this metric was then adjusted for age and sex. Cell type outliers were excluded from the dataset, and subsequent analyses were restricted to individuals of European descent. A Cox proportional-hazards model was used to evaluate the association between mtDNA-CN and time to incident neurodegenerative disease, adjusting for age and sex.

## DECLARATIONS

### Ethics approval and consent to participate

Approval access for the datasets in this study was obtained from the GTEx and UKBiobank resources.

### Consent for publication

**Not applicable**

### Availability of data and materials

RNA-sequencing data used for analyses was downloaded from the GTEx portal (http://gtexportal.org/home/datasets) on 06/18/2019 and phenotypes were obtained from dbGaP (phs000424.**v8**.p2). Aligned whole genome sequences were downloaded from google cloud. UKBiobank data was accessed under application number 17731. All in-house scripts can be found in the following Github repository: htpps://github.com/syyang93/mtDNA_GE_scripts.

### Authors’ contributions

Concept and design: Yang, Arking

Acquisition, analysis, or interpretation of data: Yang, Castellani, Longchamps, Pillalamarri, Arking Drafting manuscript: Yang, Arking

Critical revision of the manuscript: Yang, Castellani, Longchamps, Pillalamarri, O’Rourke, Guallar, Arking Obtaining funding: Arking

## Acknowledgements

This research was conducted using data from the Genotype-Tissue Expression (GTEx) project (dbGaP accession: phs000424.**v8**.p2). The GTEx project was supported by the Common Fund of the Office of the Director of the National Institutes of Health, and by NCI, NHGRI, NHLBI, NIDA, NIMH, and NINDS. This research was also conducted using the UK Biobank Resource under Application Number 17731.

