## Supplemental Information for "Blood-derived mitochondrial DNA copy number is associated with gene expression across multiple tissues and is predictive for incident neurodegenerative disease"

**Supplementary Table 1. GTEx effect size estimates between mtDNA-CN and known correlates are comparable to ARIC mtDNA-CN effect size estimates from previous literature. Power calculations are based on R^2^ values between ARIC mtDNA-CN and correlates.**

|  | **Covariate** | **GTEx  effect size estimate** | **GTEx  standard error** | **GTEx  p-value** | **ARIC  effect size estimate** | **ARIC  standard error** | **ARIC  p-value** | **Power** |
| --- | --- | --- | --- | --- | --- | --- | --- | --- |
|  | Age | -0.06 | 0.05 | 0.19 | -0.02 | 0.01 | 0.004 | 28.68% |
|  | Sex (Female) | 0.14 | 0.10 | 0.17 | 0.46 | 0.06 | 9.95e-14 | 79.44% |
|  | Neutrophils | -0.19 | 0.05 | 5e-05 | -0.14 | 0.02 | 9.97e-16 | 97.48% |

**Supplemental Table 2. Scaled median mtRNA expression in non-blood tissues does not seem to be significantly associated with blood-derived mtDNA-CN. Only Uterus and Heart – Left Ventricle pass nominal significant (p<0.05) for associations between blood-derived mtDNA-CN and tissue-specific mtRNA expression.**

|  | **Tissue** | **Effect size estimate** | **Standard error** | **P-value** |
| --- | --- | --- | --- | --- |
|  | Whole Blood | 0.15 | 0.03 | 9.10e-09 |
|  | Uterus | 0.29 | 0.09 | 0.004 |
|  | Heart - Left Ventricle | 0.12 | 0.05 | 0.017 |
|  | Skin - Sun Exposed (Lower leg) | -0.05 | 0.03 | 0.055 |
|  | Heart - Atrial Appendage | 0.07 | 0.04 | 0.084 |
|  | Artery - Tibial | 0.05 | 0.03 | 0.111 |
|  | Brain - Cerebellum | 0.07 | 0.05 | 0.130 |
|  | Esophagus - Mucosa | 0.06 | 0.04 | 0.156 |
|  | Cells - Cultured fibroblasts | -0.07 | 0.05 | 0.177 |
|  | Breast - Mammary Tissue | 0.05 | 0.04 | 0.185 |
|  | Artery - Aorta | 0.04 | 0.03 | 0.194 |
|  | Brain - Putamen (basal ganglia) | -0.05 | 0.04 | 0.207 |
|  | Brain - Cortex | 0.06 | 0.05 | 0.216 |
|  | Nerve - Tibial | 0.04 | 0.03 | 0.237 |
|  | Colon - Transverse | 0.04 | 0.04 | 0.241 |
|  | Minor Salivary Gland | 0.09 | 0.09 | 0.292 |
|  | Stomach | 0.06 | 0.06 | 0.292 |
|  | Thyroid | 0.03 | 0.03 | 0.294 |
|  | Pancreas | 0.08 | 0.08 | 0.317 |
|  | Testis | 0.05 | 0.05 | 0.318 |
|  | Esophagus - Gastroesophageal Junction | -0.04 | 0.05 | 0.427 |
|  | Brain - Nucleus accumbens (basal ganglia) | 0.04 | 0.05 | 0.443 |
|  | Brain - Caudate (basal ganglia) | 0.03 | 0.04 | 0.515 |
|  | Adrenal Gland | 0.05 | 0.07 | 0.530 |
|  | Lung | 0.02 | 0.04 | 0.566 |
|  | Esophagus - Muscularis | 0.02 | 0.04 | 0.576 |
|  | Colon - Sigmoid | 0.02 | 0.03 | 0.584 |
|  | Cells - EBV-transformed lymphocytes | 0.08 | 0.16 | 0.609 |
|  | Brain - Anterior cingulate cortex (BA24) | -0.02 | 0.05 | 0.635 |
|  | Muscle - Skeletal | -0.01 | 0.03 | 0.667 |
|  | Brain - Substantia nigra | 0.03 | 0.08 | 0.671 |
|  | Spleen | 0.02 | 0.06 | 0.671 |
|  | Pituitary | 0.02 | 0.05 | 0.673 |
|  | Brain - Amygdala | 0.03 | 0.07 | 0.704 |
|  | Vagina | 0.03 | 0.09 | 0.710 |
|  | Skin - Not Sun Exposed (Suprapubic) | 0.01 | 0.02 | 0.758 |
|  | Liver | 0.02 | 0.06 | 0.763 |
|  | Brain - Hippocampus | -0.01 | 0.05 | 0.767 |
|  | Small Intestine - Terminal Ileum | 0.01 | 0.06 | 0.798 |
|  | Brain - Cerebellar Hemisphere | -0.01 | 0.06 | 0.817 |
|  | Artery - Coronary | -0.01 | 0.04 | 0.849 |
|  | Brain - Frontal Cortex (BA9) | 0.01 | 0.05 | 0.869 |
|  | Brain - Hypothalamus | -0.01 | 0.06 | 0.873 |
|  | Adipose - Subcutaneous | 0.00 | 0.03 | 0.897 |
|  | Prostate | -0.00 | 0.06 | 0.980 |
|  | Adipose - Visceral (Omentum) | -0.00 | 0.03 | 0.993 |
|  | Brain - Spinal cord (cervical c-1) | 0.00 | 0.06 | 0.997 |
|  | Ovary | 0.00 | 0.08 | 0.999 |

**Supplemental Table 3. Genomic inflation factors for all tested tissues, represented by the median observed chi-squared value divided by the median expected chi-squared value. Bolded tissues pass the permuted genomic inflation factor cutoff, and represent tissues with significant signal.**

| **Tissue** | **Lambda** |
| --- | --- |
| **Whole Blood** | **4.71** |
| **Brain - Putamen (basal ganglia)** | **3.36** |
| **Brain - Substantia nigra** | **1.69** |
| **Colon - Transverse** | **1.61** |
| **Nerve - Tibial** | **1.56** |
| **Esophagus - Mucosa** | **1.51** |
| **Skin - Sun Exposed (Lower leg)** | **1.49** |
| **Brain - Hippocampus** | **1.48** |
| **Testis** | **1.45** |
| **Brain - Nucleus accumbens (basal ganglia)** | **1.45** |
| **Adipose - Subcutaneous** | **1.43** |
| **Colon - Sigmoid** | **1.41** |
| **Esophagus - Muscularis** | **1.41** |
| **Breast - Mammary Tissue** | **1.37** |
| **Brain - Hypothalamus** | **1.35** |
| **Skin - Not Sun Exposed (Suprapubic)** | **1.33** |
| **Muscle - Skeletal** | **1.32** |
| **Brain - Cerebellum** | **1.32** |
| **Heart - Left Ventricle** | **1.31** |
| **Vagina** | **1.30** |
| **Spleen** | **1.29** |
| **Lung** | **1.28** |
| **Pancreas** | **1.28** |
| **Adrenal Gland** | **1.27** |
| **Adipose - Visceral (Omentum)** | **1.26** |
| **Thyroid** | **1.26** |
| **Brain - Cortex** | **1.26** |
| **Esophagus - Gastroesophageal Junction** | **1.25** |
| **Heart - Atrial Appendage** | **1.24** |
| **Brain - Caudate (basal ganglia)** | **1.24** |
| **Liver** | **1.23** |
| Stomach | 1.19 |
| Brain - Spinal cord (cervical c-1) | 1.18 |
| Small Intestine - Terminal Ileum | 1.17 |
| Brain - Cerebellar Hemisphere | 1.15 |
| Artery - Tibial | 1.14 |
| Pituitary | 1.14 |
| Artery - Aorta | 1.13 |
| Brain - Frontal Cortex (BA9) | 1.12 |
| Prostate | 1.11 |
| Ovary | 1.07 |
| Minor Salivary Gland | 1.06 |
| Cells - Cultured fibroblasts | 1.03 |
| Brain - Anterior cingulate cortex (BA24) | 1.02 |
| Artery - Coronary | 1.01 |
| Brain - Amygdala | 0.973 |
| Uterus | 0.928 |
| Cells - EBV-transformed lymphocytes | 0.835 |

**Supplemental Table 4. Spearman’s rank correlation coefficients between blood estimates and specific tissue estimates for all genes that were significant in blood. Average baseline correlation is represented by the correlation for 100 randomly selected gene sets (100 genes per set). Tissues with correlations significantly different from baseline are bolded.**

|  | **Tissue** | **Spearman correlation for significant blood genes** | **Average spearman correlation for 100 permuted random gene sets** | **95% Confidence Interval** |
| --- | --- | --- | --- | --- |
|  | **Adipose - Subcutaneous** | **0.185** | **0.103** | **0.026;0.181** |
|  | **Adipose - Visceral (Omentum)** | **0.266** | **0.150** | **0.078;0.222** |
|  | **Adrenal Gland** | **0.196** | **0.090** | **0.019;0.162** |
|  | Artery - Aorta | 0.174 | 0.109 | 0.028;0.190 |
|  | **Artery - Coronary** | **-0.001** | **0.098** | **0.028;0.168** |
|  | **Artery - Tibial** | **0.177** | **0.100** | **0.026;0.174** |
|  | Brain - Amygdala | 0.074 | 0.067 | -0.014;0.148 |
|  | Brain - Anterior cingulate cortex (BA24) | 0.069 | 0.023 | -0.062;0.107 |
|  | Brain - Caudate (basal ganglia) | 0.100 | 0.061 | -0.028;0.150 |
|  | **Brain - Cerebellar Hemisphere** | **0.137** | **0.051** | **-0.026;0.127** |
|  | Brain - Cerebellum | 0.181 | 0.105 | 0.027;0.182 |
|  | **Brain - Cortex** | **0.145** | **0.050** | **-0.022;0.122** |
|  | **Brain - Frontal Cortex (BA9)** | **0.187** | **0.034** | **-0.046;0.113** |
|  | **Brain - Hippocampus** | **0.139** | **0.022** | **-0.066;0.110** |
|  | **Brain - Hypothalamus** | **0.191** | **0.058** | **-0.022;0.137** |
|  | **Brain - Nucleus accumbens (basal ganglia)** | **0.220** | **0.046** | **-0.032;0.124** |
|  | Brain - Putamen (basal ganglia) | 0.127 | 0.088 | 0.007;0.169 |
|  | Brain - Spinal cord (cervical c-1) | 0.095 | 0.094 | 0.007;0.180 |
|  | **Brain - Substantia nigra** | **0.171** | **0.015** | **-0.054;0.085** |
|  | **Breast - Mammary Tissue** | **0.318** | **0.148** | **0.080;0.215** |
|  | Cells - Cultured fibroblasts | -0.061 | -0.045 | -0.12;0.029 |
|  | Cells - EBV-transformed lymphocytes | -0.047 | 0.021 | -0.061;0.102 |
|  | Colon - Sigmoid | 0.105 | 0.143 | 0.065;0.221 |
|  | **Colon - Transverse** | **0.210** | **0.105** | **0.031;0.179** |
|  | Esophagus - Gastroesophageal Junction | 0.119 | 0.091 | 0.01;0.172 |
|  | Esophagus - Mucosa | 0.141 | 0.166 | 0.091;0.241 |
|  | Esophagus - Muscularis | 0.148 | 0.144 | 0.066;0.221 |
|  | **Heart - Atrial Appendage** | **0.203** | **0.115** | **0.035;0.194** |
|  | Heart - Left Ventricle | 0.252 | 0.189 | 0.119;0.260 |
|  | **Liver** | **0.169** | **0.076** | **-0.003;0.155** |
|  | Lung | 0.245 | 0.168 | 0.087;0.249 |
|  | Minor Salivary Gland | 0.117 | 0.105 | 0.027;0.182 |
|  | Muscle - Skeletal | 0.188 | 0.157 | 0.076;0.237 |
|  | **Nerve - Tibial** | **0.259** | **0.082** | **0.009;0.155** |
|  | **Ovary** | **0.089** | **-0.035** | **-0.112;0.041** |
|  | **Pancreas** | **0.266** | **0.097** | **0.030;0.165** |
|  | **Pituitary** | **0.258** | **0.091** | **0.012;0.170** |
|  | Prostate | 0.139 | 0.113 | 0.036;0.190 |
|  | **Skin - Not Sun Exposed (Suprapubic)** | **0.216** | **0.145** | **0.076;0.214** |
|  | **Skin - Sun Exposed (Lower leg)** | **0.066** | **0.155** | **0.083;0.227** |
|  | Small Intestine - Terminal Ileum | 0.183 | 0.162 | 0.081;0.242 |
|  | **Spleen** | **0.226** | **0.130** | **0.051;0.210** |
|  | Stomach | 0.186 | 0.144 | 0.061;0.226 |
|  | **Testis** | **0.072** | **0.153** | **0.077;0.230** |
|  | **Thyroid** | **0.055** | **0.146** | **0.065;0.227** |
|  | Uterus | 0.124 | 0.090 | 0.011;0.169 |
|  | **Vagina** | **0.167** | **0.054** | **-0.033;0.141** |

**Supplementary Figure 1. mtDNA-CN exhibits a batch effect associated with nucleic acid extraction date. The batch effect was identified using GTEx version 7 data (phs000424.v7.p2). All subsequent analyses were done with version 8, and version 8 whole genome sequences extracted prior to January 2013 were not downloaded.**

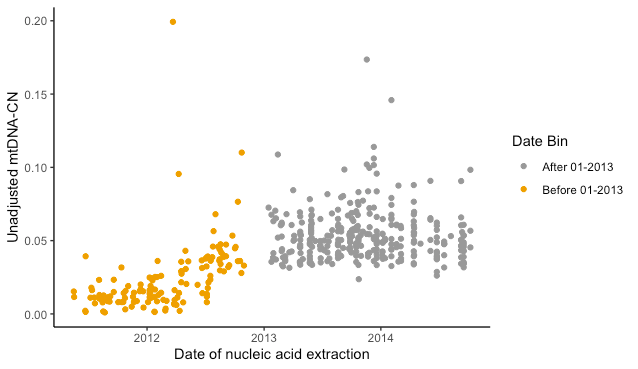

**Supplementary Figure 2. Surrogate variables 1 and 2 derived from whole blood RNA-sequencing capture cohort effects. Pairwise plots for the top 10 surrogate variables derived from whole blood RNA-sequencing, colored by cohort. Surrogate variables are protected from mtDNA-CN.**

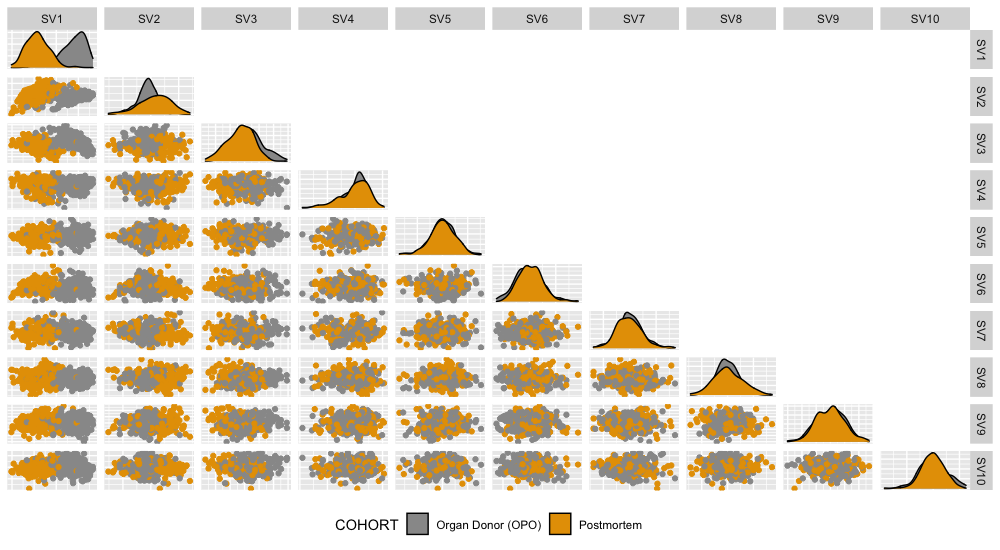

**Supplementary Figure 3. mtDNA-CN is associated with mitochondrial gene expression. Forest plot of associations between blood-derived mtDNA-CN and mtDNA-encoded gene expression in whole blood. Effect size estimates reflect the change in standard deviations for gene expression per standard deviation in mtDNA-CN.**

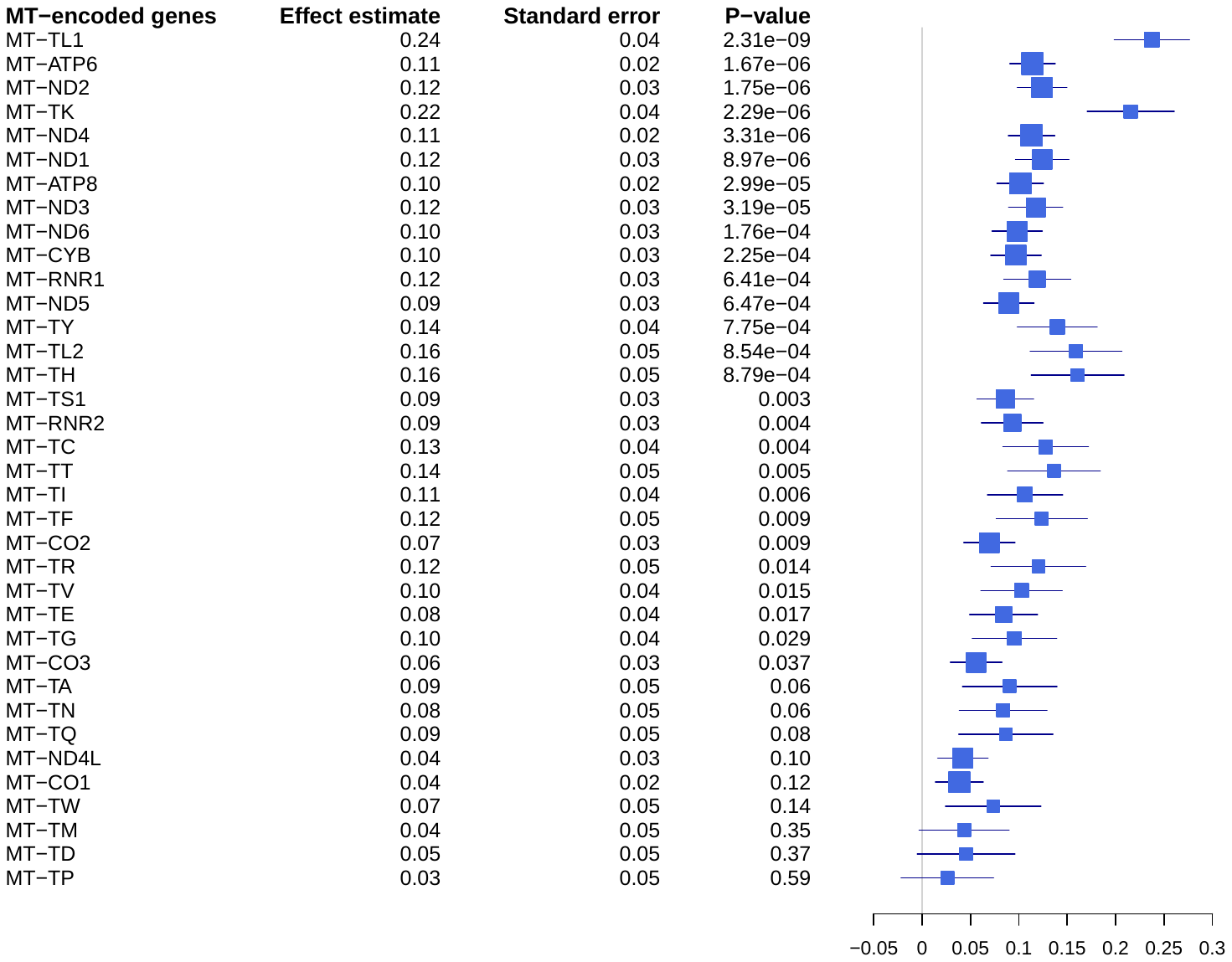

**Supplemental Figure 4. Inflation of test statistics from multiple linear regressions between gene expression in whole blood and blood-derived mtDNA-CN. Results from two-stage permutations (in gray) show no inflation of test statistics.**

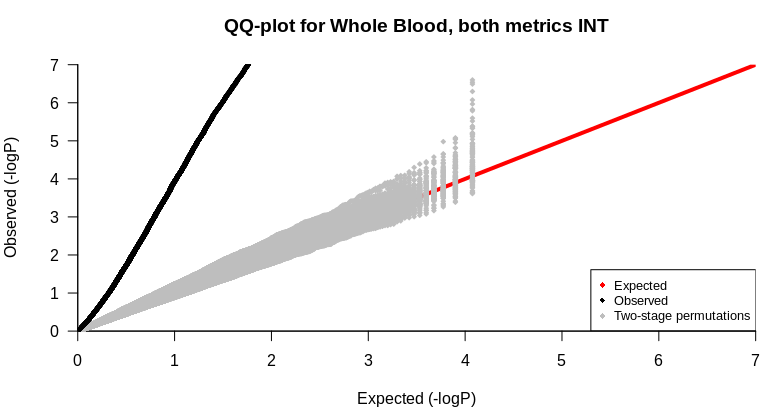

**Supplemental Figure 5. Effect size estimates between blood-derived mtDNA-CN and gene expression in blood are significantly positively shifted. Shown in gray is the percent of nominally significant genes that were positive from multiple linear regressions between blood-derived mtDNA-CN and gene expression in whole blood for permuted datasets. Dashed red line represents the actual observed value.**

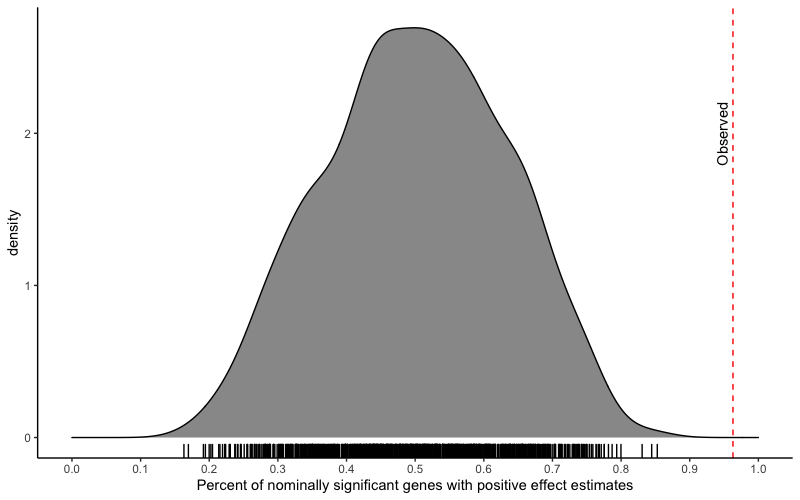

**Supplemental Figure 6. ELK1 is significantly associated with mtDNA-CN 12 out of the 18 tissues where ELK1 targets are enriched for association with mtDNA-CN. N represents number of samples for each tissue. Summary estimates calculated using a random-effects meta-analysis, excluding the effect of whole blood.**

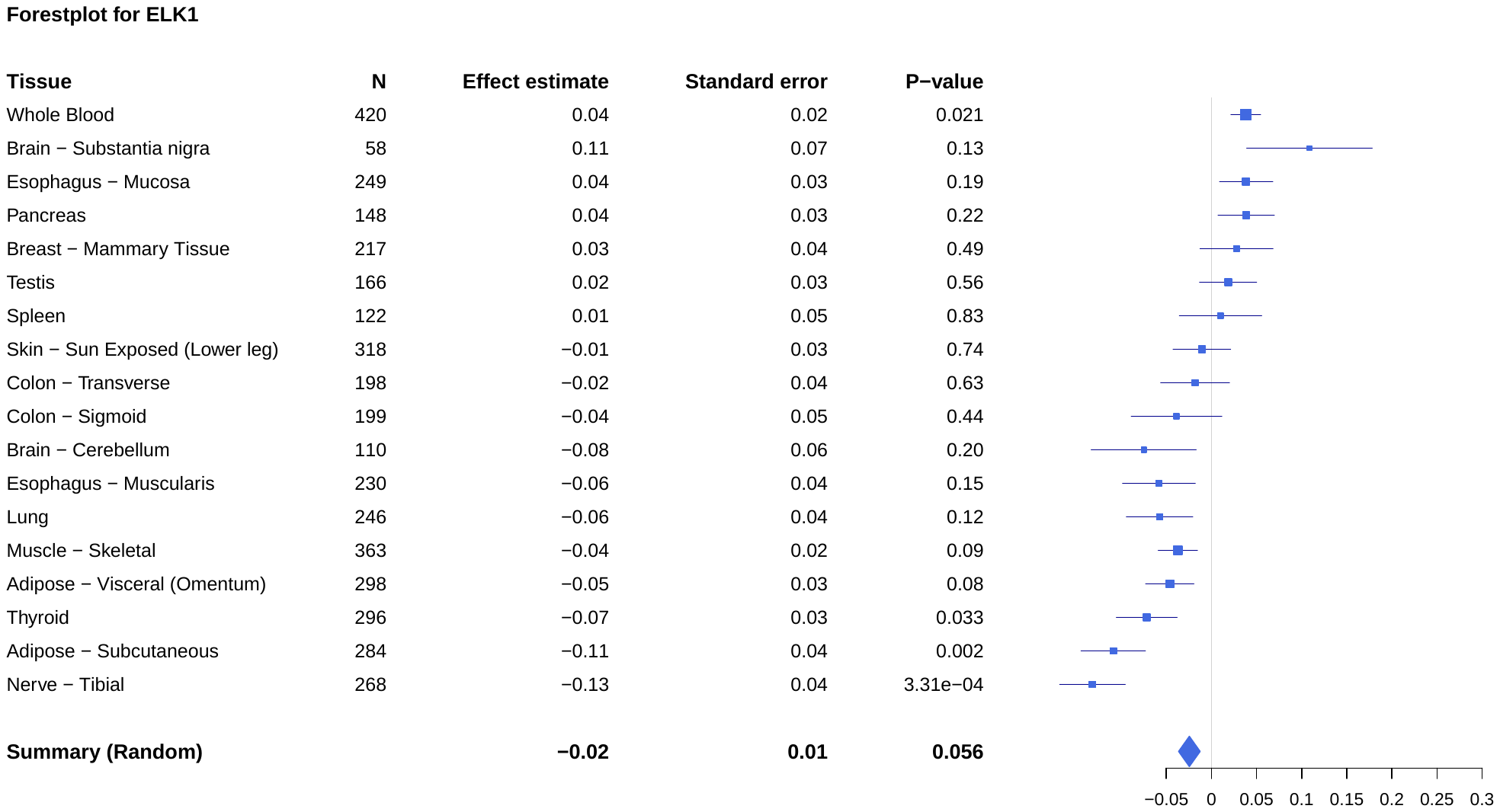

**Supplemental Figure 7. GTEx visualization showing gene expression of ELK1 in multiple tissues, downloaded on 5/9/2020 from** <https://gtexportal.org/home/gene/ELK1>.

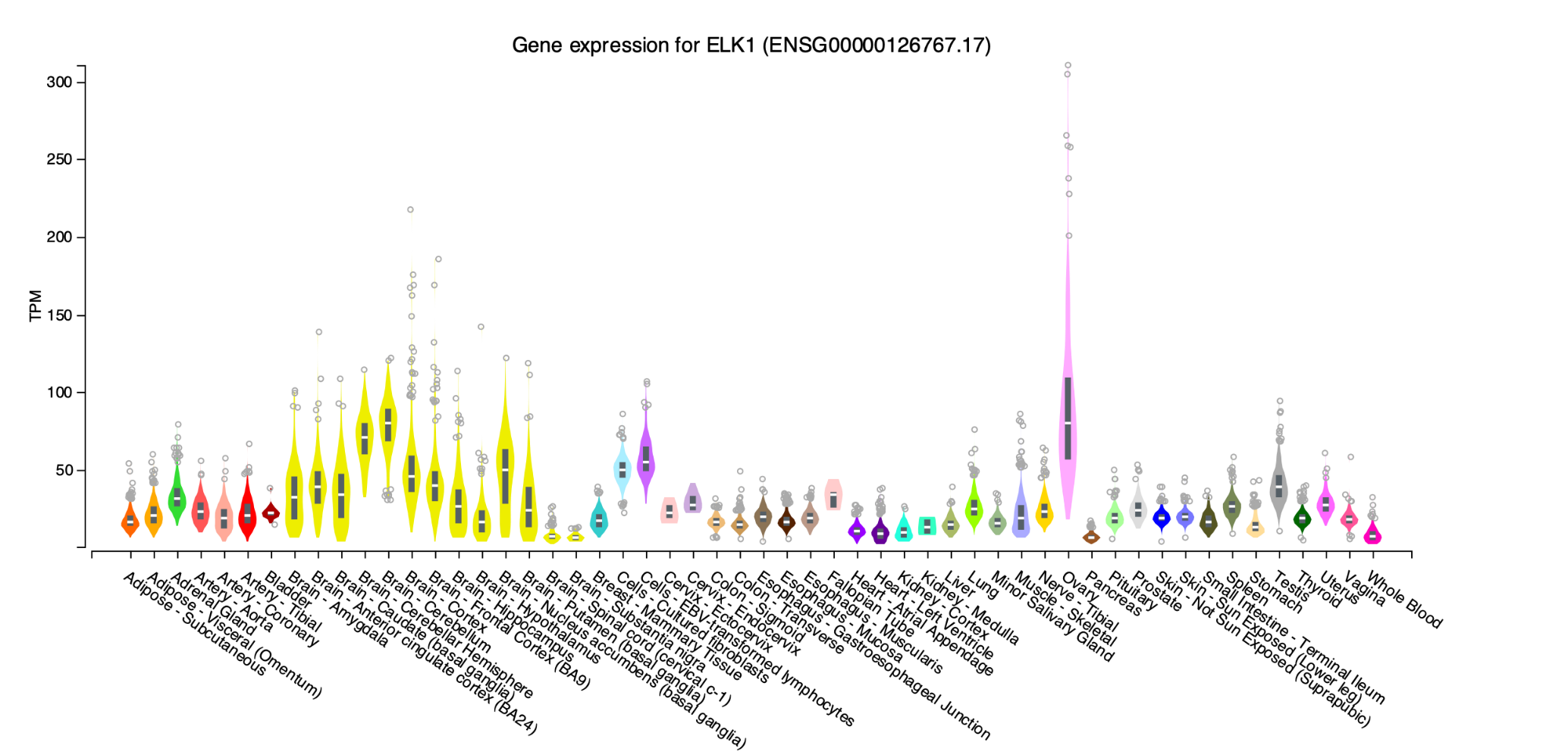

**Supplementary Figure 8. Forest plots showing t-test results for genes in neurodegenerative pathways in brain tissues. Importantly, since absolute values for t-scores were taken, negative betas do not represent significant enrichment.**

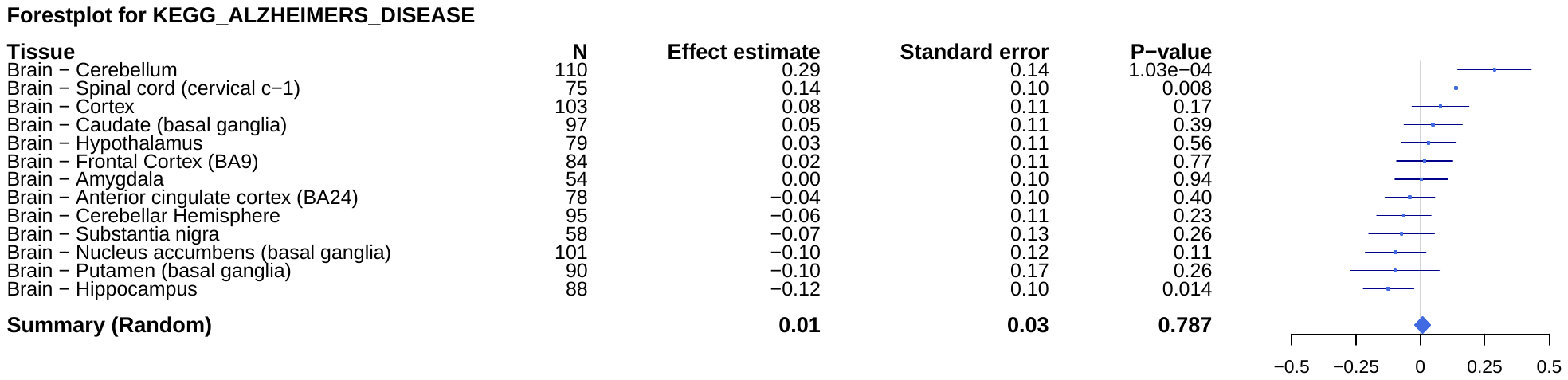

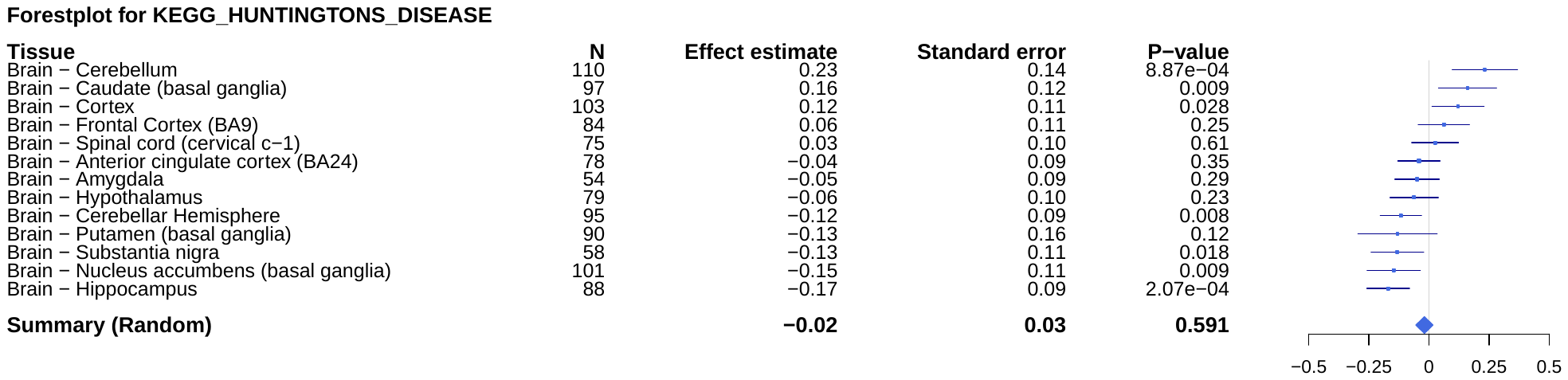

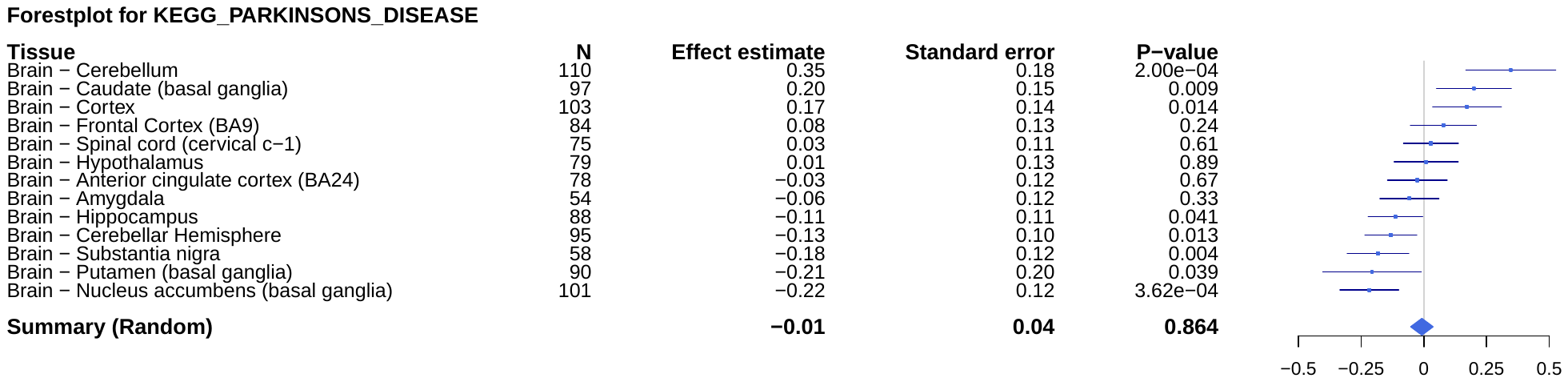

**Supplementary Figure 9. Filtering pipeline for whole blood analysis and the number of individuals retained at each step.**

Individuals with WGS data (833)

Individuals with blood RNAseq data available (629)

WGS was run post Jan-2013 (452)

Remove samples with high unaligned reads (442)

Remove cell type outliers (431)

Drop surgery individual (430)

Iteratively remove surrogate variable outliers (420)

Complete phenotype information (418)
